# Conflict detection in a sequential decision task is associated with increased cortico-subthalamic coherence and prolonged subthalamic oscillatory response in the beta band

**DOI:** 10.1101/2020.06.09.141713

**Authors:** Zita Eva Patai, Tom Foltynie, Patricia Limousin, Harith Akram, Ludvic Zrinzo, Rafal Bogacz, Vladimir Litvak

## Abstract

Making accurate decisions often involves the integration of current and past evidence. Here we examine the neural correlates of conflict and evidence integration during sequential decision making. Patients implanted with deep-brain stimulation (DBS) electrodes and age-matched healthy controls performed an expanded judgement task, in which they were free to choose how many cues to sample. Behaviourally, we found that while patients sampled numerically more cues, they were less able to integrate evidence and showed suboptimal performance. Using recordings of Magnetoencephalography (MEG) and local field potentials (LFP, in patients) in the subthalamic nucleus (STN), we found that beta oscillations signalled conflict between cues within a sequence. Following cues that differed from previous cues, beta power in the STN and cortex first decreased and then increased. Importantly, the conflict signal in the STN outlasted the cortical one, carrying over to the next cue in the sequence. Furthermore, after a conflict, there was an increase in coherence between the dorsal premotor cortex and subthalamic nucleus in the beta band. These results extend our understanding of cortico-subcortical dynamics of conflict processing, and do so in a context where evidence must be accumulated in discrete steps, much like in real life. Thus, the present work leads to a more nuanced picture of conflict monitoring systems in the brain and potential changes due to disease.

## Introduction

Whether it is deciding which method of transportation to take to get to work most efficiently or which horse to bet on to maximize monetary gain, humans are constantly integrating noisy evidence from their environment and past experience, in order to optimize their decisions. Often the information comes at intervals, thus necessitating a system that can track incoming signals over time and only commit to making a choice after sufficient evidence has been integrated (Ratcliff, 1978; Busemeyer and Townsend, 1993; Usher and McClelland, 2001), a process that has been proposed to rely on the cortico-basal-ganglia circuit (Bogacz et al., 2010). Research in human patients with implanted electrodes for clinical deep-brain stimulation (DBS) treatment has pointed to the role of the subthalamic nucleus (STN) of the basal ganglia as a decision gate-keeper. The STN is postulated to set the decision threshold in the face of conflicting information by postponing action initiation until the conflict is resolved (Frank, 2006). As predicted by the model, STN activity is increased for high conflict trials and STN-DBS affects decision making in the face of conflicting evidence (Frank et al., 2007; Coulthard et al., 2012; Green et al., 2013). Furthermore, the decision threshold correlated specifically with changes in STN theta oscillatory power (Cavanagh et al., 2011; Herz et al., 2016). Recent evidence has also pointed to the role of beta oscillations during conflict (Zavala et al., 2018). Thus, oscillatory activity, primarily in the theta and beta bands, in the basal ganglia, reflects immediate inhibition to motor output during situations involving conflict (Frank, 2006), whether it is the response, sensory or cognitive uncertainty (Bonnevie and Zaghloul, 2019).

The majority of previous studies in the STN employed paradigms in which the putative processes of conflict detection and setting of decision threshold happened in close temporal proximity. For example, in previously used paradigms such as the flanker task (Zavala et al., 2015), go-no-go (Alegre et al., 2013; Benis et al., 2014), and Stroop task (Brittain et al., 2012) evidence was presented simultaneously. Although STN activity was also studied in random dot motion paradigm that required evidence accumulation over time (Herz et al., 2018), it was unknown exactly what sensory evidence was presented when, on individual trials, due to the noisy nature of stimuli. As a result, previous studies do not allow us to fully disentangle the neural correlates of ongoing evidence accumulation and conflict during decision making. In particular, it is not clear what kind of conflicting information during evidence accumulation the STN responds to: does it respond to a local conflict, when a new piece of information does not match single previous piece in the sequence, or global conflict, when a new piece of information does not match overall evidence from the entire trial?

An important role in shaping the STN activity is played by the interaction between the cortical circuits and the STN. However, the nature and cortical locus of this interaction has only been examined in a handful of studies. Resting-state coherence between the STN and ipsilateral frontal cortex has shown a peak in the beta band in human patients (Litvak, Jha, et al., 2011; West et al., 2020) as well as rodent models of Parkinson’s disease (Magill et al., 2004; West et al., 2018). Additionally, coherence in the theta band from frontal sites (as measured with electroencephalography) to the STN increased during a conflict detection task (Zavala et al., 2014, 2016).

To precisely characterize how the neural activity in cortex and the STN changes during the process of evidence accumulation, we recorded STN local field potential (STN-LFP) simultaneously with whole-head magnetoencephalography (MEG) while Parkinson’s disease patients performed an expanded judgement task (Leimbach et al., 2018). Here, cues are presented at discrete intervals, and evidence for the correct answer develops as the participant samples and integrates multiple cues over the course of the trial (Figure 1). This paradigm allowed us to investigate how behavioural and neural responses depend on the continual unfolding of evidence extended in time, determine what kind of conflicting information the STN responds to, and test predictions of computational models.

**Figure 1:**
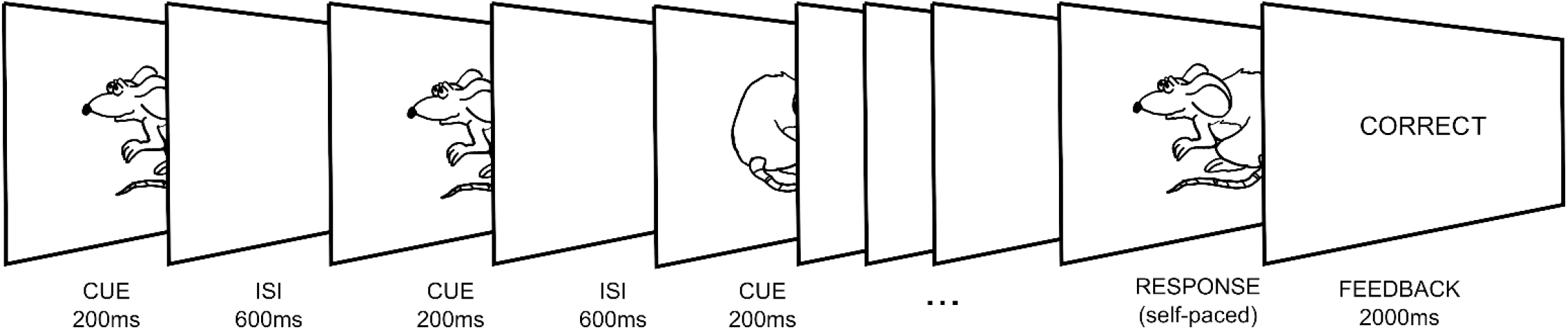
Expanded Judgement Task. Participants performed a version of an evidence integration task, with two key elements: 1. the cues were presented sequentially within the trial rather than simultaneously, which allowed us to examine evidence accumulation over time, and 2. the trial duration, i.e. number of cues sampled, was up to the participants, who responded when they felt they had received enough information to make a decision. Participants were required to guess the likely direction (left or right) the mouse ‘would run’ in. Each cue was 70% valid, i.e. they represented the correct direction 70% of the time if they were to be treated in isolation.

## Materials and Methods

### Participants

We tested 15 patients with a clinical diagnosis of Parkinson’s disease (14 male, mean age: 59, range 47-71, two left-handers), following electrode implantation for DBS treatment, before full closure of the scalp, thus allowing for intracranial recordings of the STN (all bilateral recordings, except 1 patient right unilateral and 1 patient with 3 contacts in the left STN and only 2 on the right, this patient was also subsequently diagnosed with Multiple Systems Atrophy). Among tested patients, 11 had Medtronic 3389 electrodes, while 4 had Boston Vercise™ directional leads. The surgical procedures are described in detail in (Foltynie et al., 2011). All patients were assessed on medication (mean Levodopa Equivalent Dosage 1272mg, range: 500-1727.5mg). Unified Parkinson’s Disease Rating Scale (UPDRS) part 3 scores were 39.6±14 (mean±standard deviation, range: 18-61) when OFF medication, and 15.4±6.5 (range: 7-30) when ON medication. None of the patients had cognitive impairment (Mini–Mental State Examination (MMSE) scores: mean 28.8, range: 26-30, one patient score missing), clinical depression, or apathy. Two patients were excluded from the analysis due to poor performance of the task (see *Task* below). We recruited 13 age and gender matched controls (12 male, mean age: 57, range 44-70, two left-handers). The patient study was approved by the UK National Research Ethics Service Committee for South Central Oxford and the control study was covered by University College London Ethics Committee approval for minimum risk magnetoencephalography studies of healthy human cognition. All participants gave written informed consent. Patients did not receive financial compensation and the controls were compensated for their time according to our centre’s standard hourly rate.

### Surgical Procedure

Bilateral DBS implantation was performed under general anaesthesia using a stereotactic (Leksell frame G, Elekta) MRI-guided and MRI-verified approach without microelectrode recording as detailed in previous publications (Holl et al., 2010; Foltynie et al., 2011). Two stereotactic, preimplantation scans were acquired, as part of the surgical procedure, to guide lead implantation; a T2-weighted axial scan (partial brain coverage around the STN) with voxel size of 1.0×1.0 mm^2^ (slice thickness=2 mm) and a T1-weighted 3D-MPRAGE scan with a (1.5 mm)^3^ voxel size on a 1.5T Siemens Espree interventional MRI scanner. Three dimensional distortion correction was carried out using the scanner’s built-in module. Target for the deepest contact was selected at the level of maximal rubral diameter (∼5 mm below the AC-PC line). To maximise DBS trace within the STN, the target was often chosen 1.5 - 2 mm posterolateral to that described by Bejjani (Bejjani et al., 2000). Stereotactic imaging was repeated following lead implantation to confirm placement.

### Task

To investigate the neural basis of evidence accumulation over time, we used the expanded judgement task (Figure 1, similar to the task previously used by Leimbach et al, 2018). Participants were shown a series of images of a mouse facing either left or right. Cues were presented for 200ms, with an inter-stimulus interval (ISI) of 600ms, so there was 800ms interval from one onset to another, to which we refer as Stimulus Onset Asynchrony (SOA). Participants were required to judge in which direction the mouse will ‘run’, based on the probabilities extracted from a series of sequential cue images, and then respond accordingly. The validity of the cues was 70%, such that each cue (left or right mouse) represented the correct choice 70% of the time. The two directions were equally likely across trials, thus the chance level in the task was 50%. If the participants responded based on one of the cues only, without accumulating information over time, then their expected success rate would be 70%. Responses were made by pressing a button with the thumb of the congruent hand after a self-chosen number of cues, when the participant felt they had enough evidence to make a decision. Prior to the recording, the participants underwent a short training session where they were first asked to respond only after seeing a set number of stimuli (between two and ten) and then told that for the main experiment they will decide themselves how many stimuli to observe. This was to ensure that participants chose to respond based on accumulating evidence from a sequence of images rather than just the first stimulus. Participants performed up to 200 trials (Patients: 168±11; Controls: 200 each, except one control who completed 150 trials). Two patients were excluded from the analysis due to poor performance of the task (accuracy at chance level).

### Recording and Analysis

Participants performed the task while seated in a whole-head MEG system (CTF-VSM 275-channel scanner, Coquitlam, Canada). For patients, STN-LFP, electrooculography (EOG) and electromyography (EMG) recordings were also obtained using a battery-powered and optically isolated EEG amplifier (BrainAmp MR, Brain Products GmbH, Gilching, Germany). STN-LFP signals were recorded referenced to a common cephalic reference (right mastoid).

All preprocessing was performed in SPM12 (v. 7771, http://www.fil.ion.ucl.ac.uk/spm/, (Litvak et al., 2011b)), and spectral analysis and statistical tests were performed in Fieldtrip (http://www.ru.nl/neuroimaging/fieldtrip/ (Oostenveld et al., 2011)) using the version included in SPM12.

STN-LFP recordings were converted offline to a bipolar montage between adjacent contacts (three bipolar channels per hemisphere; 01, 12, and 23) to limit the effects of volume conduction from distant sources (for more details see Litvak et al., 2010 and Oswal et al., 2016b). Four of the patients had segmented DBS leads (Vercise™ DBS directional lead, Boston Scientific, Marlborough, USA). In these cases, we averaged offline the signals from the 3 segments of each ring and treated them as a single ring contact. Thus, for each participant, we had a total of 3 STN EEG channels in each hemisphere (except for 2 participants: one with right side electrodes only, thus 3 channels, and one with 1 contact on the right excluded due to extensive noise, thus 5 channels). The LFP data were downsampled to 300Hz and high-pass filtered at 1Hz (Butterworth 5^th^ order, zero phase filter).

A possibly problematic but unavoidable feature of our task was that the stimuli were presented at relatively short SOA not allowing for the power to return to baseline before the next stimulus was presented. Furthermore, the SOA was fixed making entrainment and anticipation possible. These were deliberate design choices to be able to collect a large number of trials for model-based analyses. Any jittering of the SOAs (which would have to go in the direction of increasing their duration) would have led to far fewer trials being collected. The total duration of the recording had to be kept short as the patients were unable to tolerate extended periods of testing. Furthermore, having a very long SOA would make it more likely that the participants would resort to explicit counting, which was something we aimed to avoid.

To account for these design issues, we developed an unconventional way of performing time-frequency analysis on these data in the absence of a baseline. We first ran time frequency analysis on continuous LFP data (multitaper method (Thomson, 1982) 400ms sliding window, in steps of 50ms) on *a priori* defined beta power (13-30 Hz average = 21.5Hz; note that when looking at individual participant beta power around the response period, we found a similar band as defined *a priori*: individual mean range: 16.6-28.4Hz; overall min: 11Hz, max: 31Hz). Separately we also estimated the power in the theta band (2-8Hz average = 5Hz, e.g. Herz et al., 2016). The resulting power time series were log-transformed and high-pass filtered at 0.5 Hz (Butterworth 5^th^ order, zero phase filter) to remove fluctuations in power that were slower than our SOA. Afterwards, the power time series were epoched around the presentation of each cue stimulus (−500 to 800ms). We averaged power across contacts within each hemisphere, resulting in 1 left and 1 right STN channel, and we also calculated the mean STN signal by combining hemispheres. We used a permutation cluster-based non-parametric test to correct for multiple comparisons across time (the duration of the whole cue epoch (0-800ms) and report effects that survive correction only (p<0.05 family-wise error (FWE) corrected at the cluster level).

Similarly to LFP, MEG data were downsampled to 300Hz, and high-pass filtered at 1Hz (Butterworth 5th order, zero phase filter). For sensor-level analysis, we used only the control group data, as in the patients the sensor signals were contaminated by ferromagnetic wire artefacts (Litvak et al., 2010).

For the MEG sensor-level time-frequency analysis, we used all channels and a frequency range of 1-45Hz. All other analyses were identical to the LFP pipeline reported above. However, we corrected for multiple comparisons across all MEG channels, timepoints (0-800ms) and frequencies (1-45Hz), and only report effects that survived that correction (p<0.05 FWE corrected at the cluster level).

For source-level analysis, the continuous MEG data were projected to source space with Linearly Constrained Maximum Variance (LCMV) beamformer (Veen et al., 1997) using a 10-fold reduced version of the SPM canonical cortical mesh (Mattout et al., 2007) as the source space (resulting in 818 vertices and the same number of source channels). The source orientation was set in the direction of maximum power. See Litvak et al., (2012) for details on beamforming and Litvak et al. (2010) for details on issues regarding beamformer use for removing artefacts from simultaneous MEG and intracranial recordings. Next, time-frequency analysis was performed on continuous source data the same way as for STN-LFP except the frequencies of interest were informed by the sensor-level analysis. This biased the statistical test for discovery of an effect (cf. double dipping, Kriegeskorte, Simmons, Bellgowan, & Baker, 2009) but our aim in this analysis was post-hoc interrogation of the effects established at the sensor level in terms of their location in the cortex rather than hypothesis testing (Gross et al., 2012). To limit our search space for the coherence analysis (below), we only investigated sources that survived p<0.05 FWE correction.

Time-resolved coherence was then computed between the identified cortical sources and STN contacts by going back to raw source time series. The data were epoched (−1000 to 1000ms to increase the window for analysis), and time-frequency analysis was performed as described above with coherence between the sources and the left and right STN also computed from the cross-spectrum. Non-parametric permutation testing between conditions was corrected for multiple comparisons across channels (source vertices), time (0-1600ms to cover both cue ‘i’ and cue ‘i+1’) and frequencies (1-30Hz), and we only report effects that survive correction (p<0.05 FWE corrected at the cluster level). For completeness, we also ran an alternative connectivity measure, debiased weighted phase lag index, which is less sensitive to unequal trial numbers across conditions and volume conduction effects.

### Reconstruction of electrode locations

We used the Lead-DBS toolbox (http://www.lead-dbs.org/ (Horn and Kühn, 2015)) to reconstruct the contact locations. Post-operative T2 and T1 images were co-registered to pre-operative T1 scan using linear registration in SPM12 (Friston et al., 2007). Pre- (and post-) operative acquisitions were spatially normalized into MNI_ICBM_2009b_NLIN_ASYM space based on preoperative T1 using the Unified Segmentation Approach as implemented in SPM12 (Ashburner and Friston, 2005). DBS electrode localizations were corrected for brain shift in postoperative acquisitions by applying a refined affine transform calculated between pre- and post-operative acquisitions that was restricted to a subcortical area of interest as implemented in the brain shift correction module of Lead-DBS software. The electrodes were then manually localized based on post-operative acquisitions using a tool in Lead-DBS specifically designed for this task. The resulting locations were verified by an expert neurosurgeon.

### Choice Strategy

In order to analyse the strategy used by the participants during choice, we investigated which factors influence commitment to a choice on a given trial. We considered two factors:

The first of them is the evidence integrated for the chosen option. Such accumulated evidence was computed from Equation 1 that continuously updates the evidence (decision variable, *DV*) for a choice at time *t* based on the existing *DV* from the previous stimuli and the new incoming stimulus *S*_*t*_, where *S* =− 1 for the left stimulus, and *S*_*t*_ = 1 for the right stimulus. At the start of each trial, the decision variable was initialized to *DV*_0_ = 0.

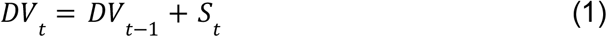

The second factor we considered was whether the stimulus was the same as the previously presented one, i.e. *SA*_*t*_ = 1 if *S*_*t*_ = *S*_*t*-1_ and *SA*_*t*_ = 0 otherwise. For all stimuli excluding the first stimulus on each trial (for which it is not possible to define *SA*_*t*_) we performed a logistic regression predicting if the choice has been made after this stimulus, i.e. we tried to predict a variable *D*_*t*_ = 1 if choice made after stimulus *t* and *D*_*t*_ = *0* otherwise. For each participant, we looked at the significance of the two factors.

### Estimating accumulated evidence using computational models

In order to analyse if STN activity reflects the amount of available evidence for each response based on the stimuli presented so far, we employed computational models that can estimate this quantity at each point in time. We compared how well different models of evidence accumulation could capture the behaviour of different patients, and then generated regressors for each patient based on the best model for that patient. In addition to the model assuming evidence is integrated according to Equation 1, we also considered three extended models which included a forgetting term (λ), a bonus term (ω), or both (Equations 2-4).

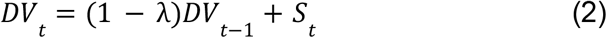

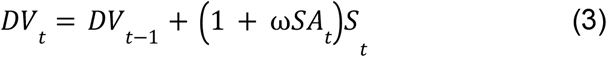

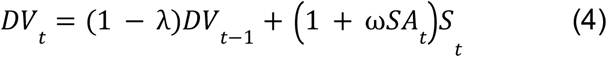

The forgetting term was used to model the decay of memory over the course of the trial and the bonus term is a weighting of ‘same’ pairs, i.e. the stimuli which match the directly preceding one (e.g.: in a ‘left-left-right’ sequence the second left stimulus would be weighted extra as it is the same as the first one).

To estimate the parameters (λ, ω), we assumed that the ratio of making a right choice to making a left choice is related to decision variable according to:

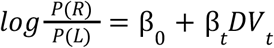

For each participant, we looked for parameters that maximized the likelihood of participant’s behaviour after all stimuli shown to that participant.

We found the winning model (based on Bayesian information criterion) to be variable across participants (number of participants in patients/control group indicated): M1 = 1/2; M2 = 0/0; M3 = 4/9; M4 = 8/2, although the model that included bonus terms was the most common.

### Estimating Bayesian normalization term

We investigated if the STN activity follows a pattern predicted by a computational model of the basal ganglia (Bogacz et al., 2007; Bogacz and Larsen, 2011). This model suggests that the basal ganglia compute the reward probabilities for selecting different actions according to Bayesian decision theory. These probabilities are updated after each stimulus and the updated information is fed back to the cortex via the thalamus. An action is initiated when the expected reward under a particular action exceeds a certain threshold. The model attributes a very specific function to the STN: ensuring that if the probability of one action goes up, the probabilities of the others go down at the same time by normalising all probabilities so that they add up to one.

In order to create regressors for neural activity recorded from the STN, we used the original proposal that the STN computes the normalization term of the Bayesian equation during the evidence integration process (Bogacz & Gurney, 2007). We defined 2 cortical integrators *Y*_*L*_ and *Y*_*R*_, which integrate evidence for the left and right stimulus respectively, as described above. Additionally, we subtracted the STN normalization term from the cortical integrators after each stimulus input in a sequence (Bogacz et al., 2016). For each participant, we assumed the integration follows one of the models described by Equations 1-4, which best describes given participants (see previous subsection). So, for example, for participants best described by Equation 1, the integrators were updated as follows

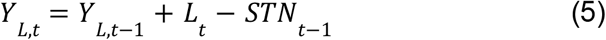

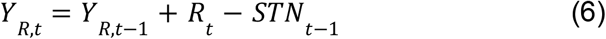

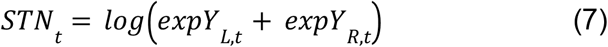

In the above equations, *L*_*t*_ = 1, *R*_*t*_ = *0* if cue *t* is left, and *L*_*t*_ = 0, *R*_*t*_ = 1, otherwise. However, for models 2-4 we added decay to the cortical integrators and bonus terms to Equations 5-6 analogously to Equation 2-4, i.e. we ensured that *DV*_*t*_ = *Y*_*R,t*_ − *Y*_*L,t*_. At the start of each trial, the integrators were initialized to *Y*_*L,*0_ = *Y*_*R,*0_ = *log*0. 5 (corresponding to equal prior probabilities of the two responses). The value computed from Equation 7 was used as Bayesian normalization regressor in Figure 2.

**Figure 2:**
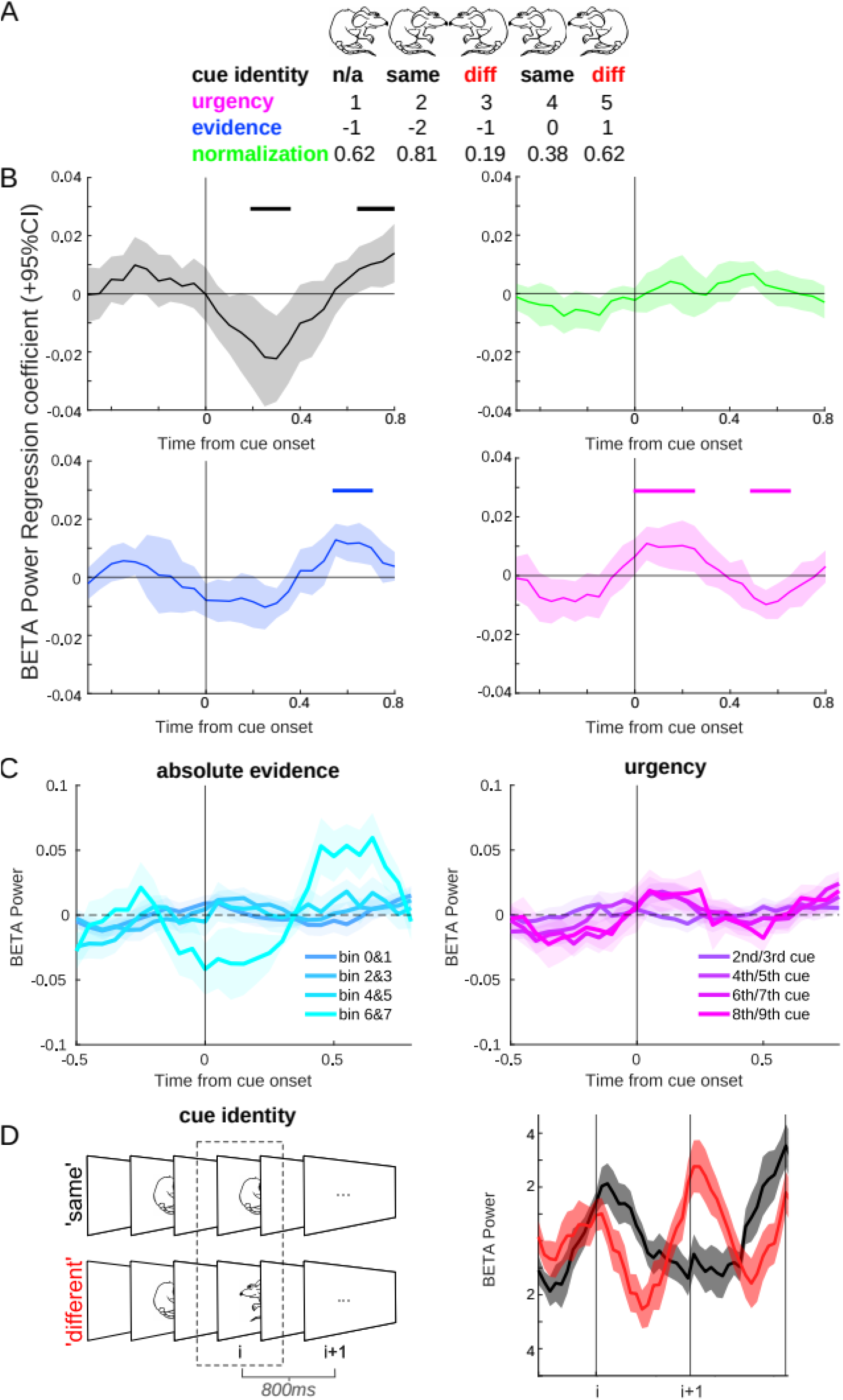
STN activity encodes local conflict and variables related to accumulation of evidence via beta oscillations. **A)** Example sequence of cues, with each regressor value shown below. For example, evidence for the ‘right’ facing mouse goes up during the first two cues, but then the appearance of a ‘left’ mouse reduces the evidence for a right response. **B)** Results of the combined GLM. A linear regression of beta power in the STN revealed that a clear signal was related to the identity of the cue (‘same’ or ‘different’, shaded in grey), absolute integrated evidence, and sample number in the sequence of cues in a trial (or ‘urgency’, i.e. the number of stimuli presented so far that could influence a general tendency to make a choice or working-memory load). Horizontal lines represent significant times after cluster correction for multiple comparisons. There was no encoding of Bayesian normalization in the STN signal, as proposed previously (Bogacz et al., 2007, 2016). Note that although the regressors are presented separately for easier visualization, they were included in a combined GLM. All regressors were z-scored before entering the model. We did not find any effects when regressing theta band activity in the STN with the above regressors. **C)** Raw beta power plotted as a function of binned evidence (left) or cue number (right), as well as for cue identity **(D)**, note this latter panel is identical to part of Figure 3B.

## Results

### Patients are able to accumulate evidence over time

Patients waited on average 6.6 stimuli before making a response (6.59±0.52 sem) and their accuracy was significantly above the 70% level expected if they only based their decision on a single cue (80±0.03 sem, t=3.6, p=0.004). Controls waited on average 6.3 stimuli before making a response (6.29±0.46 sem) and were similarly above 70% in their accuracy (88.6±0.01 sem, t=18.4, p<0.001). There was no significant difference between groups in the number of stimuli viewed before making a choice (t=0.42, p-value = 0.68), but patients had lower accuracy (t=-2.99, p=0.0009) and slower reaction time (as measured from the onset of the last cue before a response was made, t=2.16, p=0.041). See Table 1 for summary of behavioural measures.

**Table 1:**
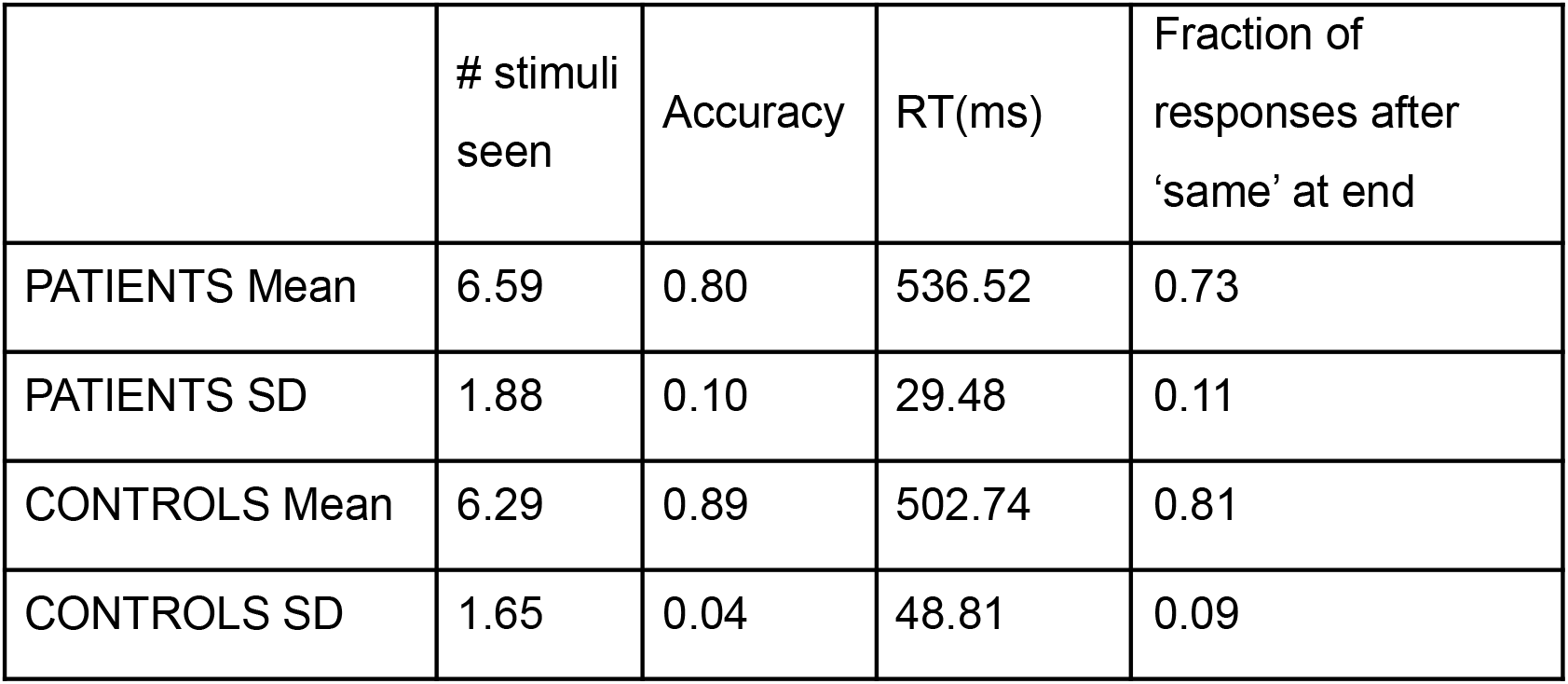
Behavioural results showing mean and standard deviations for each group. RT: Reaction time; ACC: accuracy. The analytical probability of a ‘same’ pair at the end of the sequence would be 58% if participants chose the moment of response randomly. Both patients and controls responded significantly more often after a ‘same’ pair (both groups p<0.001).

To explore potential strategies participants could have used in the task, we compared performance in both groups to an agent that would have been an optimal observer, and would choose to respond left if the number of left cues was higher than the number of right cues, to respond right for a larger number of right cues, and would choose randomly if the numbers were equal. In other words, for each participant, we calculated the accuracy they would have achieved had they integrated evidence optimally, having seen the stimuli sampled by the participant on each trial. We found that controls and patients had significantly lower accuracy (controls: p=0.019, patients: p=0.0076) than an ideal observer would have, based on the same cue sampling (89% for controls and 87% for patients).

Next, we asked whether participants were just solving the task by responding after they spotted two of the same stimuli in a row (i.e. after the first ‘same’ pair). To address this question, we investigated to what extent participants’ response after stimulus was predicted by accumulated evidence, and by same stimuli in a row (see Materials and Methods for details). Most participants had responses best predicted either by accumulated evidence alone (6 patients and 6 controls), or by both accumulated evidence and stimulus repetition (5 patients and 7 controls). For remaining 2 patients none of these factors was predicting their response. Hence, there was no participant who exclusively relied of making a choice after seeing the ‘same’ stimulus, without considering evidence integrated so far.

### STN beta power reflects multiple variables related to ongoing decision making

In order to understand the impact of different variables related to the decision making process on activity in the STN, we created a combined GLM, including four regressors: *cue identity, normalization model, accumulated evidence* and *sample number*. These are described in detail below.

Cue identity was taken as a measure of ‘local conflict’, by taking all cues (excluding the first and last cues in a sequence) and categorizing them as the ‘same’ or ‘different’ from the previous cue (Figure 2A & 2D). We found that beta power carried information about the similarity of the stimulus to the previous one (‘cue identity’, 200-350 and 650-800ms, p=0.024 and p=0.032, see Figure 2B & 2D).

In addition to local conflict, we analyzed whether other variables occurring in theoretical models of decision making were reflected in neural activity. We explored if STN represents the normalization term in Bayes theorem as proposed in a previously suggested computational model (Bogacz et al., 2007). This model predicts that the activity in the STN is proportional to a logarithm of the normalization term in Bayes theorem ln P(cue i). This probability is computed on the basis of all previous cues {cue 1, …, cue i-1} so it expresses how expected the current cue is given all cues seen before. The negative of this regressor, −ln P(cue i), is equal to Shannon’s surprise, so it expresses how much cue i disagrees with overall information in all previous cues, and hence it could be viewed as a measure of global conflict. Therefore, a possible correlation between the normalization term ln P(cue i) and LFP activity could be explained by either of two hypotheses. A computational model (Bogacz et al., 2007) predicts a positive correlation, whereas a hypothesis that STN responds to global conflict predicts a negative correlation. We tested if the normalization term affects power of beta oscillations in the STN and did not find evidence supporting any of these two hypotheses in our data (Figure 2B).

We also explored whether there was a signal reflecting the magnitude of accumulated evidence in the STN. Additionally, we included a regressor on beta power equal to the serial position of the cue stimulus within a trial. Including this regressor was motivated by two observations: reports of decreasing beta power as a result of increasing working memory load (Zavala et al., 2017), and presence of “urgency signals” in the basal ganglia that increase within a trial and reflect the growing urgency to making a choice (Thura & Cisek, 2017). We found a significant effect in both regressors (absolute evidence: 550-700ms, p=0.008; cue number or urgency: 0-250 and 500-650ms, p=0.01 and p=0.02).

We did not find a clear relationship between behaviour on the task and these neural effects (see Extended Data Table 2-1). However, cue identity (early peak) showed a relationship with both RT (r=0.62,p=0.024; note if an outlier of the STN data is taken out then the correlation is no longer significant, p=0.12; outlier detected as more than 1.5 interquartile range above the upper quartile or below the lower quartile, which is appropriate when data is not normally distributed), as well as a trend for the number of cues sampled (r=0.53,p=0.064).

### STN beta power shows persistent activity to local conflict during evidence accumulation

Complementing, and extending on the above regression analyses, in order to further investigate how the STN represents the inconsistencies when faced with conflicting evidence over time, we separated all cues into two categories: ‘same’ or ‘different’ to the one immediately before it (we term this ‘cue i’, Figure 3A). In our analyses of neural responses to cues, we excluded the first cues in a sequence, because it is not possible to classify them as ‘same’ or ‘different’, and last cues seen as they overlapped with the response period. Thus, if a participant experienced this sequence of mouse images: ‘left-right-left-left-right’, the analysed conditions would be ‘different-different-same’.

**Figure 3:**
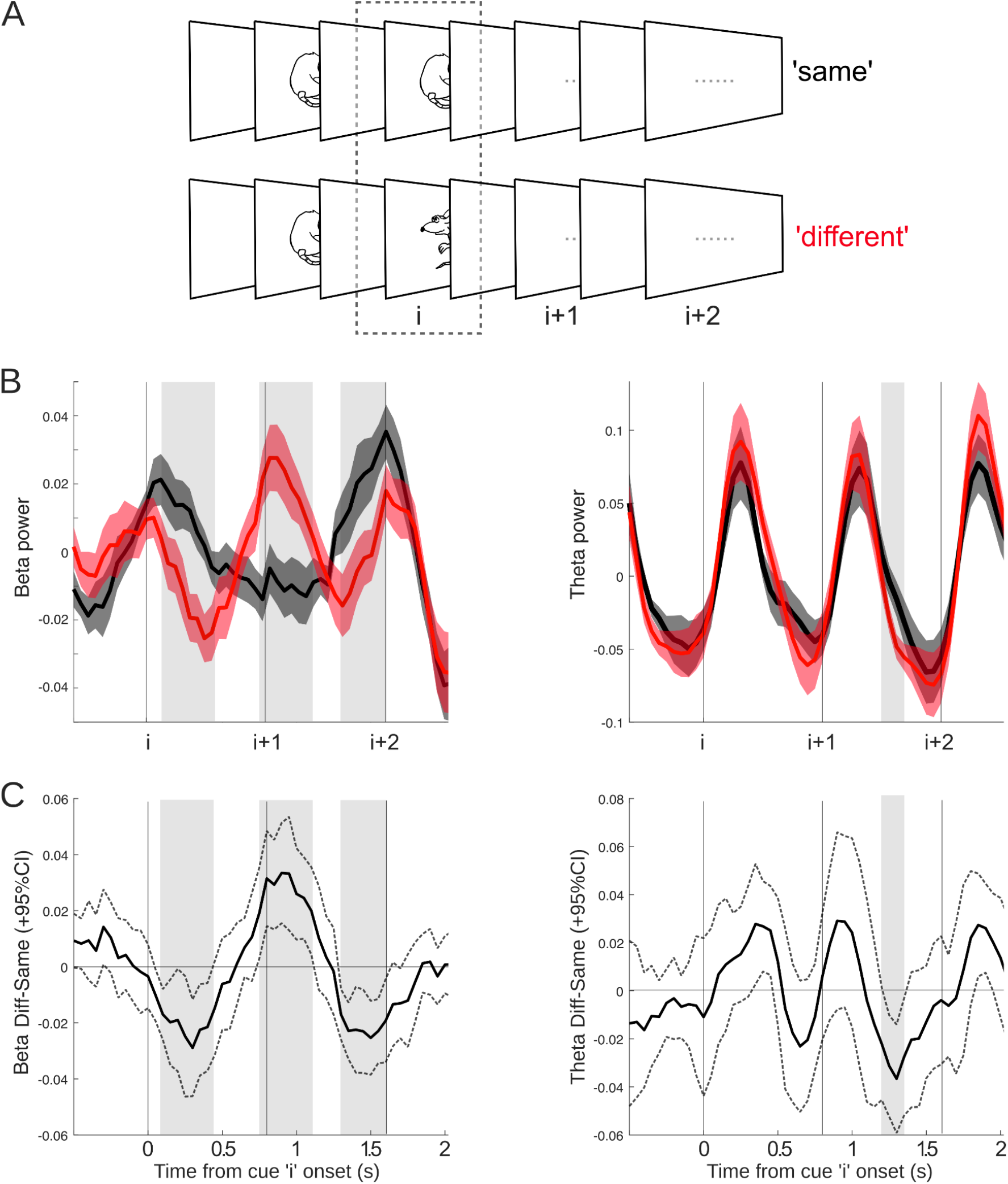
Beta signalled local conflict, and carried this effect over to the next cue in a sequence. **A)** Notation used in the paper. Let us consider an arbitrary cue i in a sequence, where i>1: If cue i-1 is the same as cue i, then we would call this the ‘same’ condition, and ‘different’ otherwise. We also plot the subsequent cues (i+1, i+2) for carry-over effects, but these are collapsed across cue type, left or right. See Extended Data Figure 3-1 for more details. **(B)** Left panel: Beta carried information locally as well as over to the next cue, with increased beta power for the ‘different’ condition. Right panel: Theta only carried mismatch information at the next cue in the sequence. Significant time periods are highlighted with shaded grey bars. Vertical lines show onset of cues in the sequence. The shaded error bars show standard error of the mean. **C)** Difference waves of conditions (‘different’ minus ‘same’) with 95% confidence intervals shown by the dotted lines. After an initial dip there is a clear increase in beta power following the conflicting cue (i) starting just before the onset of cue i+1. Significant time periods are highlighted with shaded grey bars copied from panel B for comparison. Note that the apparent onset of the effect before zero is due to limited time resolution of the time-frequency decomposition.

We found that beta oscillations (i.e raw beta power) responded to local conflict, generating a significant difference between ‘same’ and ‘different’ cues (cue ‘i’ in Figure 3B left panel) starting around 100ms after cue onset. Beta also showed a significant difference in the subsequent cue (i+1), with ‘different’ cues showing an increase in beta power, thus conflicting information on cue i results in increased beta power on cue i+1 (see Figure 3C), a pattern of activity that is consistent with response inhibition. Significant time clusters: 100-450ms (p=0.022, d=1.74), 750-1100ms (p=0.014, d=1.73), 1300-1600ms (p=0.012, d=2.40). These effects were greatly reduced in the theta band, with an effect of condition only briefly detectable during cue ‘i+1’ (Figure 3B-C, right panel).

### Cortical activity reflects rapid but non-persistent local conflict detection

We investigated sensor-level MEG signals from controls in response to local conflict detection within the sequence. As with the STN, widespread activity over central sensors was found to signal local conflict – with an initial dip followed by an increase in beta power on ‘different’ trials (Figure 4A). The dip and increase in beta power were associated with different clusters of electrodes. The first cluster showed a significant decrease to different cues in the beta band across central, and predominantly right occipital, parietal and temporal sensors (inset in Figure 4A, 0-450ms, 8-35Hz, p=0.002, Cohen’s d=1.22;). A subsequent second cluster, more restricted to central sensors, showed an increase in beta power to different cues (550-800ms, 9-25Hz, p= 0.008, Cohen’s d=1.35).

**Figure 4:**
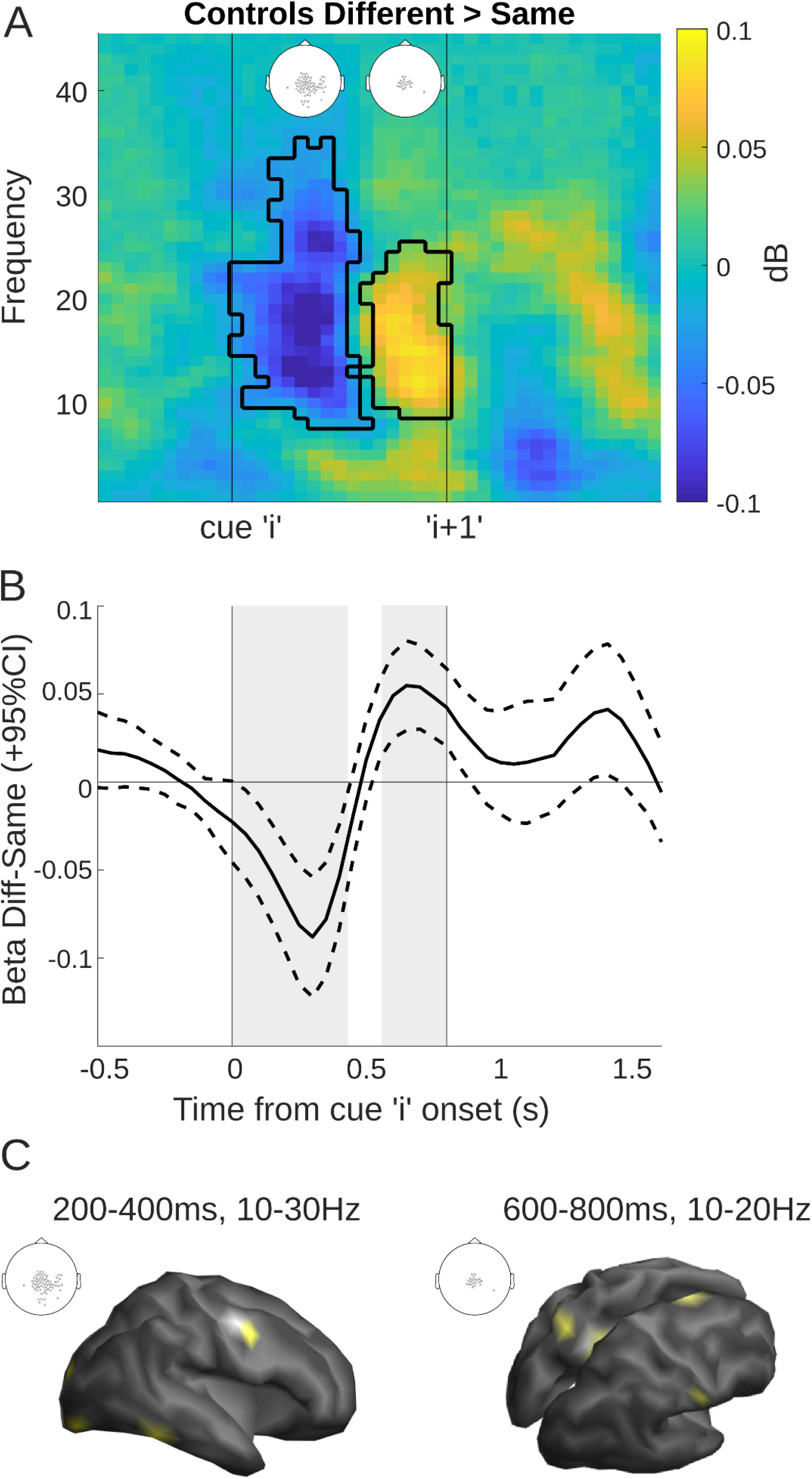
Cortical activity to local conflict parallels STN but peaks earlier on average and has a shorter time course. **A)** Time-frequency plot showing significant times and frequencies when contrasting ‘different’ vs ‘same’ cues, averaged over all significant sensors. Significant sensors are shown as an inset, separately for the 2 clusters (cluster 1: 0-450ms, 8-35Hz; cluster 2: 550-800ms, 9-25Hz,). **B)** Difference wave for the beta effects over clusters (13-30Hz) band, as represented in Figure 3B. The dotted lines indicate 95% confidence intervals. **C)** Left: Source localization in a combined sample of patients and controls revealed the source of cluster 1 in three right-lateralized areas: occipital pole, ventral temporal cortex and lateral premotor cortex (BA6). Right: Cluster 2 showed left lateralized superior parietal lobe (BA7), left posterior cingulate cortex (BA23), right primary sensory cortex and right dorsal premotor cortex/pre-supplementary motor area (dPM/BA6).

Interestingly, the time-course of the cortical effect was quicker than that of the STN (Figure 4B vs 3B), with conflicting information only lasting until the onset of the next cue in the sequence.

### Coherence is increased between STN and frontal cortex during local conflict

We used beamforming in a combined sample of patients and controls to localize the source of the ‘same-different’ effect (cluster 1: averaged over: 200-400ms [to exclude the time the stimulus was displayed on the screen], 10-30Hz; cluster 2: averaged over 600-800ms, 10-20Hz). In cluster 1 we found 3 right-hemisphere lateralized peaks (Figure 4C): occipital pole (2 peaks: MNI 19, −98, −14; 35, −89, −16), ventral temporal cortex (2 peaks: MNI 59, −53, −21; 52, −51, −21) and lateral premotor cortex (BA6, 2 peaks: MNI 52, −7, 44; 51, 3, 40). Cluster 2 was localized to left superior parietal lobe (SPL/BA7, MNI −23, −61, 52), left posterior cingulate cortex (PCC/BA23, MNI −14, −47, 31), right dorsal premotor area (dorsal/medial BA6, MNI 7, 2, 69) and right primary somatosensory cortex (BA1, MNI 61, −18, 31). Note, at an uncorrected threshold (p<0.001) we also found the lateral premotor cortex, occipital pole and temporal cortex as in cluster 1, which is expected given the overlapping topography of sensors in the two clusters.

Next, we measured in patients the coherence between these cortical vertices and both the left and right STN-LFPs, separately. The coherence spectra were averaged over adjacent vertices resulting in three cortical sources for cluster 1 and four sources for cluster 2. We found a significant increase in coherence between the right dorsal premotor cortex and the right STN (510-900ms, 10-13Hz, p=0.03, Cohen’s d=1.71; 900-1240ms, 18-24Hz, p=0.01, Cohen’s d=1.44; see Figure 5), suggesting that ipsilateral cortical-subthalamic coherence is increased in the face of local conflict in the right hemisphere. Furthermore, it seems there are two separate points of coherence over the course of the cue, one after the onset of the conflict cue and one that extends into the processing of the next cue in the sequence, this latter effect is in the mid-high beta band, possibly reflecting response inhibition. No other sources, nor the left STN showed any significant effects. For completeness based on previous reports, we also investigated coherence with the inferior frontal gyrus (which was present as a source in patients at an uncorrected threshold), and found that it did not show any significant coherence with the STN. We also used debiased weighted phase lag index as an alternative measure and found the same effects, albeit with reduced significance (cluster 1: 690-910ms,10-13Hz, p=0.043; cluster 2: 860-1150ms, 20-24Hz p=0.056).

**Figure 5:**
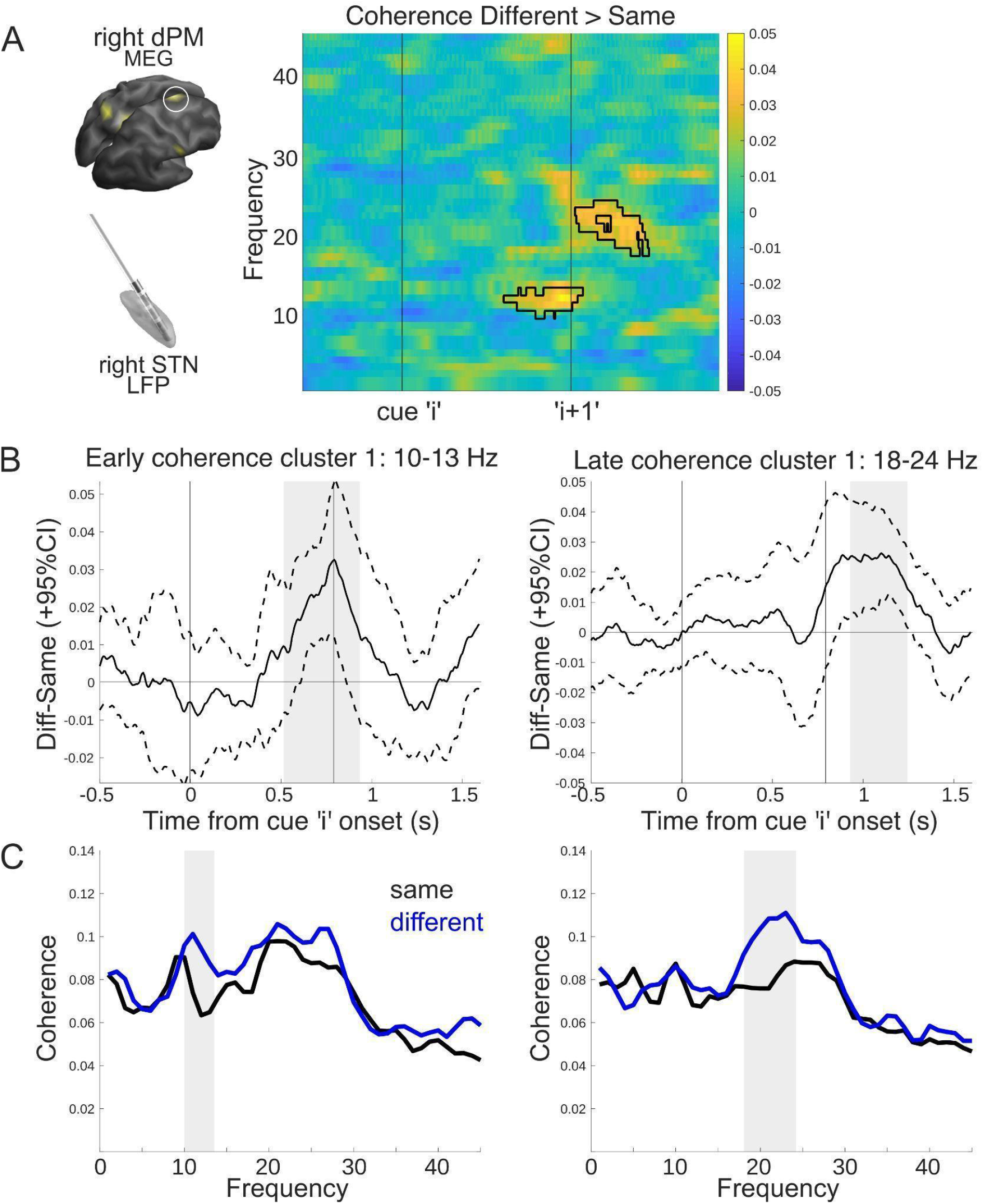
Increased coherence between right frontal cortex and right STN during local conflict. **A)** Time-frequency plot of coherence between the right STN and the right dorsal premotor cortex (visualized on the left). Two coherent clusters emerged, with an alpha/low beta coherence increase after ‘different’ cues, and a later increase in beta coherence carrying over into the next cue in the sequence. Significant clusters are shown in black outline. Inset on top left shows the source of the cortical effect for reference. **B)** Time-courses of coherence for both alpha/low and high beta plotted as a difference wave between conditions. The dotted lines indicate 95% confidence intervals. Significant timepoints are highlighted in grey. **C)** Frequency spectra of ‘same’ (black) and ‘different’ (blue) trials during the significant time period from A. Grey area highlights significant frequencies:10-13, 18-24 Hz.

## Discussion

In this experiment we present novel evidence pertaining to the role of the STN and cortico-subthalamic communication during sequential decision making, using a task in which participants had to integrate evidence over discrete time periods, with no constraints on how many samples they could observe before making a decision. We find evidence for persistent local conflict representation in the STN via beta oscillations, and increased coherence with frontal cortex. We also observed modulation of beta power in STN by evidence accumulation and number of cues presented so far in a trial.

### Representation of Conflict in the STN

We found that activity in the beta band carried information about local conflict, i.e. a difference between the current cue and the preceding one, but not about global conflict i.e. a surprise by the current cue given all previous cues. Although we established that beta power varies depending on whether the current cue differs from a previous one in a sequence – an event to which we refer as a local conflict – it is less clear from our data what the function of this activity is, and what fundamental variable it encodes.

It is possible that the observed changes in beta power are connected with motor inhibition. Beta power was initially lower for cues that were ‘different’ to the one immediately before and continued to increase across the next cue in the sequence. Activity in the beta band has been shown to carry conflict information across trials (Zavala et al., 2018), but we also show this effect within a trial, as conflict arises within the sequence of evidence. Hence, one can interpret the increase of beta power as a stop signal, or a break on motor output (Alegre et al., 2013) inhibiting a response after an inconsistent cue. Moreover, the majority of trials ended on a ‘same’ cue (Table 1), which is in line with an overall increase in beta synchronization after ‘different’ cues and lower probability of responding.

The response to different cues could also be interpreted as encoding of expectancy valuation, uncertainty or surprise. Beta power increases have been reported when a ‘surprise’ stimulus is presented (Wessel et al., 2016), and STN activity measured with fMRI has been shown to increase when there is increased uncertainty which option is correct arising due to too much choice (Keuken et al., 2015). However, in our study we found no evidence that the STN encodes the Shannon’s surprise term.

### Interaction between STN and Cortex

Interestingly, the ‘same’-’different’ effect on average peaked earlier in the cortex, and also did not carry over to the next cue in the sequence (Figure 4A). A possible interpretation is that the cortex signalled the immediate local conflict to STN, dovetailing with recent evidence suggesting the cortical conflict signal precedes the STN (Chen et al., 2020), which then maintains a more persistent activity to inhibit responses (Brittain et al., 2012; Fife et al., 2017).

When we localized the sources of the ‘same’-’different’ effect, we found the local conflict signal in widespread areas of the cortex. Only one frontal source, located in dorsal premotor cortex/supplementary motor area (dPM/BA6) showed a significant coherence modulation with the ipsilateral STN only, namely an increase in alpha/low-beta coherence shortly after the offset of a ‘different’, or conflict, cue, and an increase in beta coherence that carried over to the next cue in the sequence (Figure 5). The right BA6, specifically dorsal BA6 (Mattia et al., 2012; Mirabella, 2014), is well-established as a cortical region involved in response-inhibition/initiation and cognitive control (Chambers et al., 2007; Simmonds et al., 2008; Aron, 2011).

While it is well-established that the cortex communicates with the STN via two anatomically defined pathways, the indirect and the hyperdirect pathways (Albin et al., 1989; DeLong, 1990; Nambu et al., 2002), recent evidence suggests the existence of two separate coherent beta oscillatory networks between the cortex and the STN (Oswal et al., 2016a). Here we find evidence for two different bands of oscillatory connectivity between the STN and dorsal premotor cortex, which may have implications for understanding the involvement of various pathways in sequential evidence accumulation. Interestingly, a recent study showed evidence of a hyperdirect pathway from inferior frontal gyrus (IFG) to the STN operating in the 13-30Hz range (Chen et al., 2020), which points to a more ventral portion of the frontal cortex than presented here. In fact, many studies in stop-signal/go-nogo tasks point to the IFG (Aron et al., 2014), however in these tasks conflict is not part of an evidence accumulation process, hence we may expect differences depending on the type of decision being made, (Erika-Florence et al., 2014; Hampshire, 2015; Mosley et al., 2020).

Due to the evoked-activity as a result of the ongoing cue presentation, we were unable to reliably estimate the directionality of coherence, but previous reports on resting-state data have shown cortex to drive STN activity (Litvak et al., 2011a), which is in line with the finding here that the ‘same’-’different’ effect seems to peak earlier in the cortical signal. However, recent data has also suggested that during processing of incongruent stimuli, STN to primary motor effective connectivity is increased in the beta band (Wessel et al., 2019), suggesting that the directionality of communication may be different across task and non-task contexts.

### Where is the theta conflict signal?

The predominant theory of STN function, and also that of the cortex during conflict detection, is the involvement of theta oscillations (Cavanagh and Frank, 2014). A large portion of empirical findings on the STN shows that it carries conflict information via the theta band (Cavanagh et al., 2011; Bastin et al., 2014; Zavala et al., 2015, 2016, 2017, 2018; Herz et al., 2016). Yet in our task we only found a weak effect of theta modulation, in the cue following a local conflict (cue i+1). This effect was present only in the STN, and no theta effects were found in the cortex. Moreover, this manifested as reduced theta synchronization to ‘different’ cues, which is the opposite of the standard reported theta increase during conflict. One explanation may be the task design, as it differs from previous paradigms: there are no long intervals over which to examine slow oscillations, such as theta. Our results, therefore, though focussed on theta power, may be dominated by evoked potentials, as cues were presented in a fixed, relatively short duration sequence. Additionally, here conflict is defined over the course of multiple cues, not on a singular trial in isolation. Thus, the integration of conflict over time may in fact be driven by different signals – beta may represent a more consistent inhibition. Nevertheless, others have also reported a lack of theta effects in the STN during a stop-signal task (Bastin et al., 2014).

### Updating models of the STN

An influential model of the role of the STN in decision making proposed by Frank (2006) suggests that in situations of conflict between competing responses an increased activity of STN postpones action initiation (Frank, 2006). This model proposes that STN is essential for decision making since it ensures that an action is only selected when it has high evidence, relative to the other options. Another model proposed by Bogacz & Gurney (2007) suggests that the basal ganglia compute the reward probabilities for selecting different actions according to Bayesian decision theory (Bogacz et al., 2007; Bogacz and Larsen, 2011). While in our task we did not find conclusive evidence that the STN is encoding Bayesian normalization (Figure 2B), it is important to remember that, despite being on medication, these experiments were performed in patients whose neural circuitry has been affected by advanced Parkinson’s disease. Thus, one cannot rule out the possibility that the Bayesian normalization is encoded by the STN of healthy individuals, but testing this hypothesis would require a different experimental technique (e.g. recording of STN neural activity from animals during an analogous decision making task, such as in Brunton, Botvinick, & Brody, 2013). Evidence also suggests that subdivisions within the STN may be responsible for different types of inhibition, with prepotent response inhibition to cues (go-no-go task) being more dependent on the ventral portion of the STN (Hershey et al., 2010). Given that the majority of our recording sites were well within the dorsal (‘motor’) region of the STN, we cannot rule out the contribution of more ventral sites to these computations.

We conclude that contrary to the emphasis on theta signals in the context of immediate conflict, here we find a prominent role for beta oscillations in signalling local conflict in a sequence of evidence. We find that both frontal cortex and the STN carry this signal, and show increased coherence in the beta band that carries over to the next cue in the sequence. Thus, we show increased communication in these areas may reduce the probability of responding in the face of incoming conflicting information.

## Data availability

The full MEG dataset for controls is available in BIDS format on https://openneuro.org/datasets/ds002908 and LFP and source data for patients is available on https://data.mrc.ox.ac.uk/data-set/human-lfp-recordings-stn-during-sequential-conflict-task. Code and analysis pipeline at https://github.com/zits69/MOUSE_LFPMEG.

## Acknowledgements & Funding

This work has been supported by MRC grants MC_UU_12024/5, MC_UU_00003/1 and BBSRC grant BB/S006338/1. The Wellcome Centre for Human Neuroimaging is supported by core funding from Wellcome (203147/Z/16/Z). UK MEG community is supported by UK MEG Partnership award from the Medical Research Council (MR/K005464/1). We thank the patients for their participation. We thank Dr. Ashwini Oswal, Dr. Simon Little, Dr. David Pedrosa, Dr. Damian Herz and Dr. Viswas Dayal for clinical support of patient recordings. We are also grateful to Dr. Tim West, Dr. Hayriye Cagnan and Dr. Simon Farmer for commenting on the manuscript.

This research was funded in whole, or in part, by the Wellcome Trust [203147/Z/16/Z]. For the purpose of open access, the author has applied a CC BY public copyright licence to any Author Accepted Manuscript version arising from this submission.

